# Best Practices for Interpretable Machine Learning in Computational Biology

**DOI:** 10.1101/2022.10.28.513978

**Authors:** Valerie Chen, Muyu Yang, Wenbo Cui, Joon Sik Kim, Ameet Talwalkar, Jian Ma

## Abstract

Advances in machine learning (ML) have enabled the development of next-generation prediction models for complex computational biology problems. These developments have spurred the use of interpretable machine learning (IML) to unveil fundamental biological insights through data-driven knowledge discovery. However, in general, standards and guidelines for IML usage in computational biology have not been well-characterized, representing a major gap toward fully realizing the potential of IML. Here, we introduce a workflow on the best practices for using IML methods to perform knowledge discovery which covers verification strategies that bridge data, prediction model, and explanation. We outline a workflow incorporating these verification strategies to increase an IML method’s accountability, reliability, and generalizability. We contextualize our proposed workflow in a series of widely applicable computational biology problems. Together, we provide an extensive workflow with important principles for the appropriate use of IML in computational biology, paving the way for a better mechanistic understanding of ML models and advancing the ability to discover novel biological phenomena.

## Introduction

Machine learning (ML) has transformed the field of computational biology. The ability to acquire large-scale datasets in a high-throughput manner along with the rapid increase in computational power have enabled the development of powerful prediction models. The prediction model provides scientists the ability to query and study biological phenomenon. For this reason, a prominent goal has increasingly become to perform *knowledge discovery* by leveraging what the prediction model has learned to generate new hypotheses about unknown biological mechanisms (Eraslan et al., 2019). These hypotheses can then be evaluated in laboratory settings to guide experimentation.

However, prediction models are often complex and difficult to understand. For example, the many weights of a neural network are not immediately meaningful to a human. To address this difficulty, the field of Interpretable Machine Learning (IML) has become extremely relevant to the goal of knowledge discovery. IML methods provide additional information that can *explain* a prediction model’s score or classification, allowing scientists to gain insight into the model behavior (Miller, 2019; Doshi-Velez and Kim, 2017). However, IML methods have only begun permeating computational biology research, as reviewed in Azodi et al. (2020); Eraslan et al. (2019); Talukder et al. (2021); Novakovsky et al. (2022). There is still limited guidance on how to use these methods to unveil potential insights in a wide variety of biological contexts.

In order to use explanations to identify important hypotheses from prediction models for downstream, resource-intensive wet-lab experiments, it is imperative to establish systematic guidance. In this work, we discuss how to address two oft-overlooked, yet strong assumptions made when performing knowledge discovery using IML methods:

1. The prediction model generalizes to new data points, ensuring that the model can be used on a broad range of datasets.
2. The way in which an IML method is used to unveil hypotheses about biological mechanisms might lead to successful discoveries, ensuring that the method can guide resource-intensive experiments.

To begin building a workflow, we discuss steps in existing literature that address the above assumptions. We found that there is significant work surrounding **Model Verification** to address Assumption 1, where various approaches (e.g., *in silico* mutagenesis, external dataset evaluation) have been developed to verify the model’s ability to generalize to new data. However, explanations are commonly utilized without verifying Assumption 2 to understand limitations of the selected IML method. In limited settings, there has been some work on **Knowledge Verification** where the validity of hypotheses suggested by an IML method is compared to well-studied biological phenomenon. In this work, we adapt existing techniques from a large body of work in the IML literature (including but not limited to Adebayo et al. (2020); Kim et al. (2022); Zhou et al. (2022); Yang and Kim (2019); Chen et al. (2022a)) to introduce a more widely accessible step to verify Assumption 2 called **Explanation Verification**, which tests hypotheses unveiled by explanations against synthetic data with known ground truth. We showcase how **Explanation Verification** can be applied to a wide range of IML methods for three different computational biology applications which leverage neural networks as the prediction model. We focus our discussion on applying the verification steps to prediction models that are neural networks, because they are often considered to be the least interpretable, but the same procedure can also be applied to other types of prediction models. The paper is organized as follows. First, we provide a short background on IML, focusing on the two broad types of IML methods shown in **Fig**. 1. Next, we overview existing work on **Model Verification** and **Knowledge Verification** and introduce the **Explanation Verification** step. We then present results from performing the**Explanation Verification** step on both types of IML methods across three computational biology applications: transcription factor binding site prediction, cellular image classification, and cell phenotype prediction from gene expression data by biologically informed neural networks. Finally, we discuss open challenges to generalizing the verification steps to further applications.

**Figure 1:**
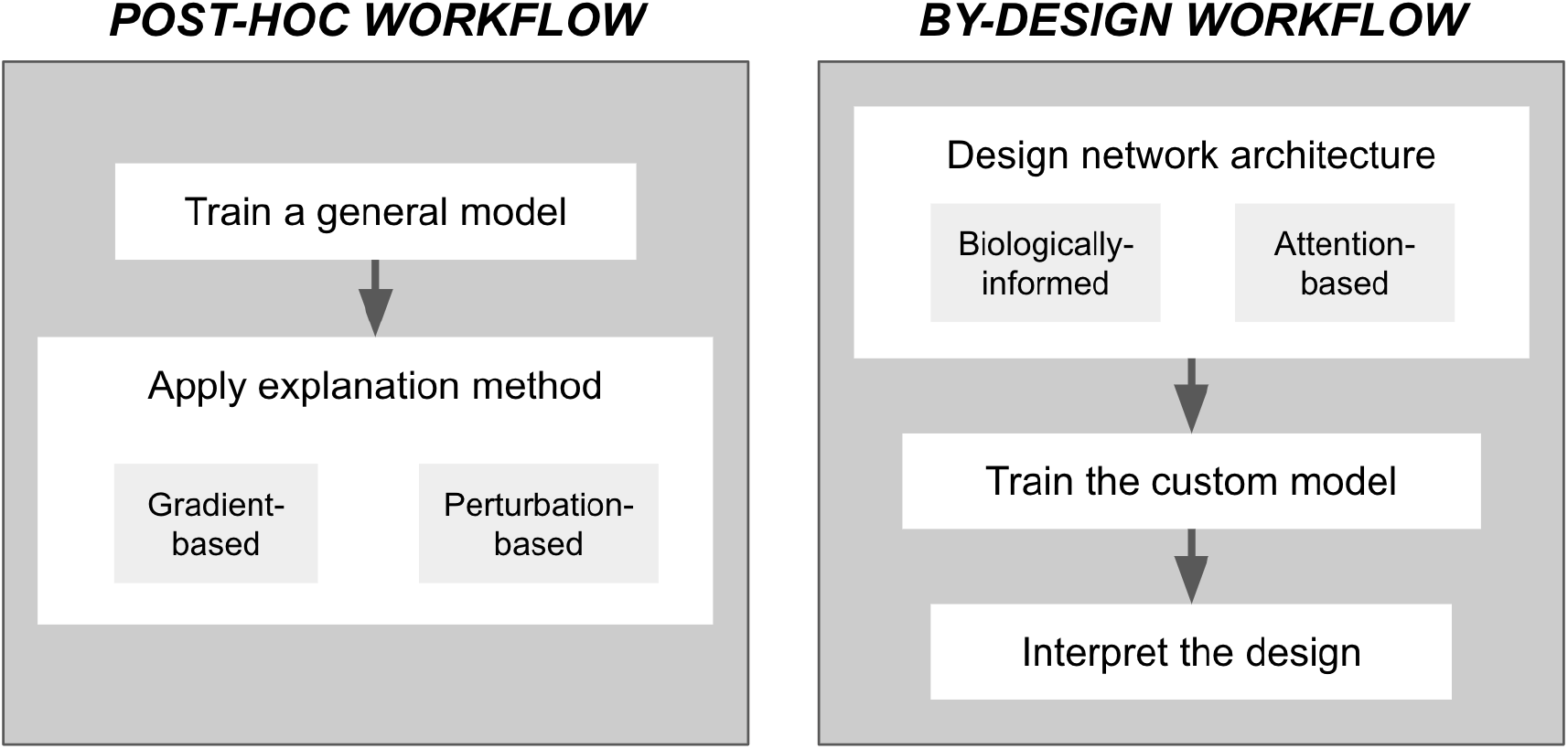
The two main IML approaches used to explain prediction models are *post-hoc* explanations and *by-design* explanations. Each approach has its own canonical workflows and popular types of IML methods.

## Preliminaries on IML Methods

Before diving into our proposed workflow, we first overview the two main IML approaches that are used to explain prediction models in computational biology applications (as shown in **Fig**. 1).

### Post-hoc Explanations

The most commonly used IML methods are post-hoc explanations, which are flexible and generally model-agnostic due to the fact that they are applied after the design and training of a prediction model. *Feature importance* methods are commonly used in computational biology applications (Novakovsky et al., 2022; Yang and Ma, 2022). These are methods that assign each input feature (e.g., a pixel in an cellular image or a DNA sequence feature) an importance value based on its contribution to the model prediction. As such, a large magnitude feature importance score would imply a large contribution. These importance values can be calculated using one of two ways: (1) gradient-based methods (e.g., DeepLIFT (Shrikumar et al., 2017), Integrated Gradients (Sundararajan et al., 2017), GradCAM Selvaraju et al. (2017)) and (2) perturbation-based methods (e.g., *in silico* mutagenesis, SHAP/DeepExplainer (Lundberg and Lee, 2017), LIME (Ribeiro et al., 2016), Fourier-based attributions (Tseng et al., 2020)). For a more detailed discussion on additional types of IML methods, we refer the readers to Chen et al. (2022b); Azodi et al. (2020).

### By-design Methods

Interpretable by-design models, as the name suggests, are models that can naturally be interpreted (Rudin, 2019). For example, a linear model is thought to be interpretable because one can easily inspect the coefficient weights to gauge the importance of each feature to the prediction outcome. Similarly, decision trees are interpretable because one can inspect the splits in the tree. Other interpretable by-design models include logistic regression, decision rules, and GAMs (Hastie and Tibshirani, 1987). While the aforementioned models are canonical by-design models in the IML literature, new by-design IML approaches that leverage recent advances and significant performance of deep neural networks are burgeoning in computational biology, either by constructing biologically-informed neural networks or by incorporating attention mechanisms.

#### Biologically-informed neural networks

are model architectures that encode domain knowledge. The process of architecture design is application-specific and, on its own, constitutes a difficult, open problem that is beyond the scope of this work. While such networks may appear to be more interpretable, the workflow and associated verification steps (introduced in later section) is crucial to ensure that the domain knowledge is being utilized by the model in the way that it was intended. Examples of biologically-informed neural networks include Ma et al. (2018), which represent the hierarchical cell subsystems capturing interwoven intracellular components and cellular processes, in the neural network design, Elmarakeby et al. (2021), which leverages the organization of experimentally-validated biological pathways, and Fortelny and Bock (2020), which incorporates the gene regulatory pathways and protein signaling pathways into the network architecture. In the above examples, the hidden nodes in the neural network are associated with biological entities, such as genes and pathways. The relative importance of the biological entities are most often evaluated by some self-defined measure, including the Relative Local Improvement in Predictive Power score defined in Ma et al. (2018) and the model weight-based importance score defined in Fortelny and Bock (2020). One could still apply a post-hoc explanation to the by-design neural networks (e.g., as in Elmarakeby et al. (2021)) to explain the model prediction using importance scores over input features rather than model internals. In addition, a recent method, named PAUSE (Janizek et al., 2022), demonstrates one way to bridge post-hoc and by-design methods. Specifically, PAUSE is a biologically-constrained autoencoder model that is explained using a post-hoc game theoretic approach.

#### Attention

is a technique that has become a popular addition to neural networks where a network incorporates and learns a set of weights which indicate how much the model is paying attention to particular part of the input (Vaswani et al., 2017). These weights do not incorporate domain knowledge, but are automatically learned as part of the training process and has been shown empirically to help the network focus on the correct parts of the input. The weights are often considered as an explanation (e.g., as in Song et al. (2021); Karbalayghareh et al. (2022); Tao et al. (2021)), although the validity of such an explanation remains debatable (Serrano and Smith, 2019).

## Methods – A Workflow on How to Use IML for Knowledge Discovery

### What is Knowledge Discovery?

The process of *knowledge discovery* using IML methods is to apply a selected IML method to a prediction model to generate new hypotheses and unveil insights about biological mechanisms. For example, hypotheses that one might be interested in gleaning from using IML methods for the three applications we showcase later include: What is the binding motif for different transcription factors? One might hypothesize motifs by selecting DNA subsequences that the explanation attributes high feature importance (Koo and Ploenzke, 2020; Avsec et al., 2021). What features indicate different cell cycle stages? One might hypothesize the important feature by selecting the most salient parts of the explanation (Nagao et al., 2020). What pathways are used in various biological processes? One might hypothesize the relevant pathways by choosing the nodes of a network with the highest weights (Fortelny and Bock, 2020). Eventually, each of these hypotheses and predictions should be further studied either by wet-lab experiments or by comprehensive comparisons with orthogonal data to support the findings. To identify the most promising explanations for further study, there are two important assumptions that should be considered when applying IML methods to reveal new biological insights:

1. The model that is being explained generalizes to new data points. If Assumption 1 is not satisfied, then knowledge discovery cannot be performed on new data points.
2. The way in which an IML method is used to unveil hypotheses about biological mechanisms might lead to successful discoveries. If Assumption 2 is not satisfied, then hypotheses identified from explanations for knowledge discovery may be misleading.

We next present verification steps as depicted in **Fig**. 2 to help address the above assumptions. The workflow includes verification steps found in the existing literature, **Model Verification** and **Knowledge Verification**, as well as a new step **Explanation Verification** to address limitations of existing verification steps. Together, the workflow will help researchers gain confidence in the hypotheses they are drawing when using IML methods.

**Figure 2:**
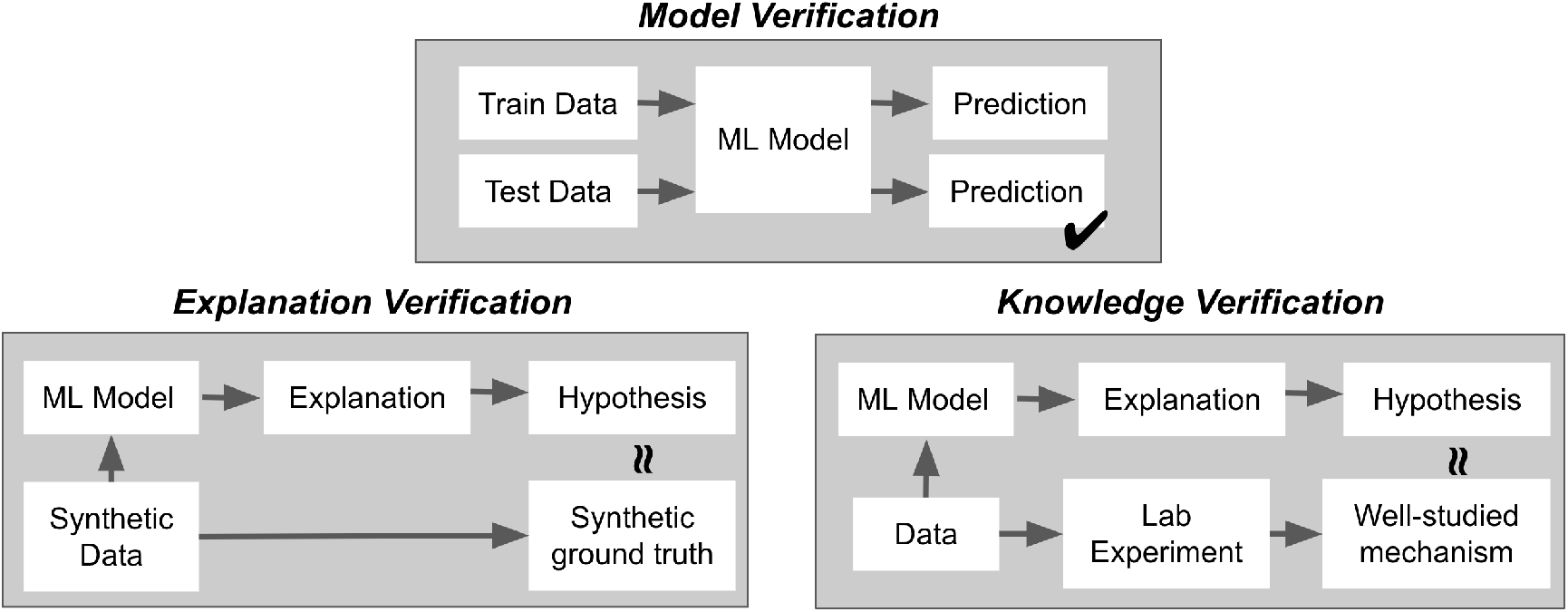
Visualizing the three steps of the proposed workflow to aid in identifying better hypotheses for knowledge discovery: **Model Verification** ensures that Assumption 1 is satisfied which is that the prediction model generalizes to new data points; **Explanation Verification** ensures hypotheses derived from explanations align with synthetic data and is particularly useful in settings where there is a lack of well-studied mechanisms; **Knowledge Verification** ensures that hypotheses from explanations align with data for which there are well-studied mechanisms. Both **Explanation Verification** and **Knowledge Verification** help address Assumption 2.

### Existing Verification Steps

To aid in selecting the best hypotheses, two existing verification steps are used to bring awareness of the capabilities and limitations of the prediction model and explanations. We first overview these existing practices and then describe why these verification steps may not be sufficient to address both assumptions.

#### Model Verification

ensures that Assumption 1 holds before even applying IML methods, meaning that the model *generalizes* to the type of data that one would be performing knowledge discovery on. If one wants to discover mechanisms on the data coming from the *same* distribution that the prediction model was trained on, then model verification can be performed by testing the model performance on a held-out validation set (discussed extensively in Heil et al. (2021)). If one wants to discover new biological phenomena on more general data, then testing the prediction model on data collected from an entirely different experimental setting should be considered (as in Coudray et al. (2018); Shao et al. (2021)). In addition, one can also introduce perturbations to existing samples and check if the prediction model accurately reproduces experimentally-validated outcomes of perturbations (as in Zhou and Troyanskaya (2015); Zhou et al. (2018); Fudenberg et al. (2020)).

#### Knowledge Verification

tests hypotheses suggested by IML methods against well-studied, prior knowledge. Note that this step is only applicable in settings where there exists well-studied prior knowledge about the biological mechanism of interest and, ideally, that mechanism would be similar to the type of knowledge that one would like to discover. This is commonly done by training a model with data for which the underlying biological mechanism is well-studied and evaluating the hypotheses suggested by the explanation against the expected behavior. For example, one might compare the features attributed by the explanation with the highest importance scores against well-studied mechanisms that have been found experimentally. **Supplementary Note** A highlights how this verification step can be performed for the three applications discussed in this work. While knowledge verification can ensure that Assumption 2 holds, there are practical limitations of performing this verification step when there is a lack of ground truth knowledge to compare explanations against, motivating our proposed verification step which we describe next.

### Introducing the Explanation Verification Step

We introduce the **Explanation Verification** step to help ensure that Assumption 2 holds in settings where there is not well-studied mechanisms and ground truth knowledge is not easily accessible.* This step verifies hypotheses from IML methods through the use of *synthetic* data to control the logic in the data, and then evaluate whether hypotheses drawn from the candidate explanation reflect the intended logic. In the next step, we demonstrate how to instantiate this step for three different applications. Similar evaluations of IML methods using synthetic data is common in the IML literature (e.g., in Adebayo et al. (2020); Kim et al. (2022); Zhou et al. (2022); Yang and Kim (2019); Chen et al. (2022a)). A synthetic dataset can also be constructed using simulators of well-studied or semi-realistic data generating distributions (Sun et al., 2021). While synthetic data may not capture all the complexities of data reflecting real biological processes, well-designed synthetic data that captures key pieces of logic is already a helpful starting point to study IML methods. Furthermore, if an explanation exhibits undesirable properties on simple synthetic data, then this step would serve as a warning to take those findings into account in following steps. Note that the use of synthetic data allows for one to test multiple variations of similar inputs, providing a quantitative measure of how consistently the explanation unveils the intended hypotheses. This quantitative measure should be noted in the explanation verification process and taken into account in when performing knowledge discovery using IML methods. Specifically, an inconsistent explanation may suggest the need to present a measure of uncertainty over the final conclusions.

## Results – Applying Explanation Verification

We instantiate the **Explanation Verification** step on three applications to illustrate potential limitations in IML methods as shown in **Fig**. 3. We use the same datasets, model architectures, and IML methods used in the original publications and follow the same approaches used to draw hypotheses from IML methods.

**Figure 3:**
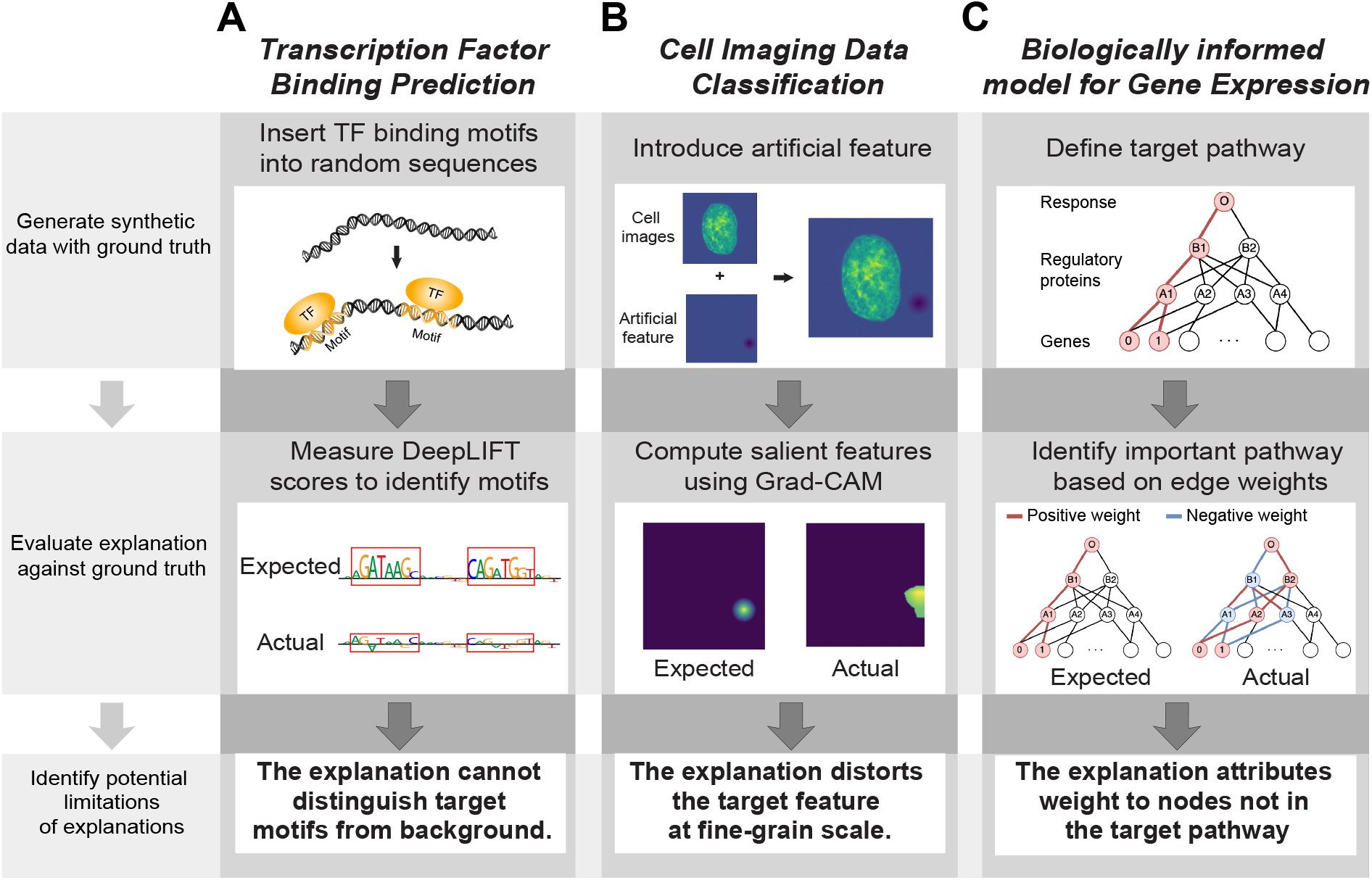
We overview the results of applying **Explanation Verification** to aid in identifying potential limitations of an IML method before using the explanation method to perform knowledge discovery. We present results for three different applications: (A) transcription factor binding site prediction, (B) cellular imaging classification, and (C) gene expression-based phenotype prediction by biologically informed neural networks. For each application, we walk through the verification’s steps (shown on the left) using prediction models and IML methods considered in prior work. In this figure, we illustrate how explanation verification can help identify potential pitfalls for each application that one should be cautious of when applying IML methods for knowledge discovery. In the text, we provide a more comprehensive discussion of each application on when explanations are helpful versus faulty.

### Transcription factor (TF) binding site prediction

TFs are regulatory proteins that bind to specific DNA sequences by recognizing the sequence patterns, called binding motifs. Deep learning models have been extensively applied in predicting TF binding sites based on DNA sequence, and IML methods have been leveraged to identify the binding motifs (Koo and Ploenzke, 2020). To showcase explanation verification, we utilize the example of BPNet (Avsec et al., 2021), which is trained on high-resolution ChIP–nexus data to predict base-resolution TF binding, and the same IML method, DeepLIFT (Shrikumar et al., 2017), to interpret the BPNet. DeepLIFT is used to identify transcription factor binding motifs by selecting clusters of DNA sequences that are attributed the highest feature importance. Here, we test when this approach to posing hypotheses may be most promising using synthetic set-ups. We sample five motifs from Kheradpour and Kellis (2014) to create multiple synthetic datasets each reflecting a different binding behavior. We study both simple binding prediction and more complex binding behaviors that include cooperative and competitive binding (Ibarra et al., 2020). An example of a complex binding behavior is shown in the top panel of **Fig**. 3 (A). In both the simple and the competitive setting, we found that DeepLIFT more consistently attributes the target motif(s) importance scores that are significantly larger than the average scores attributed to the background noise (on average 53.64% of the time across the five motifs considered), suggesting that the correct hypothesis would be identified for a majority of the time. However, in the cooperative setting, the importance scores attributed to the target motifs are more consistently indistinguishable from background noise (on average 1.51% of the time across the 20 motif pairs), meaning the correct hypothesis would not be discovered for the majority of the time. An example where the expected motif importance is high but the actual motif importance is low is shown in the middle panel of **Fig**. 3 (A). The variability in results across binding behaviors suggests the importance of this verification step to quantify an explanation’s ability to satisfy Assumption 2 under varying settings. All experimental details can be found in **Supplementary Note** B.

### Imaging Data Classification

Cellular and tissue imaging has been extensively utilized to visualize the subcellular structures within the cells or cellular patterns in tissues. Deep learning models have contributed significantly to the analysis and interpretation of such bioimaging data (Lundervold and Lundervold, 2019). Here, we illustrate how explanation verification can be adapted to cellular imaging data for the goal of classifying cell cycle phases used by Nagao et al. (2020), which trained a CNN model on real fluorescence cellular images. Nagao et al. (2020) used GradCAM (Selvaraju et al., 2017) to highlight the important regions for the prediction, hypothesizing that these key features would help to distinguish different cell phases. We construct a synthetic dataset to test whether the most salient features highlighted by GradCAM aligns with ground truth. To establish a ground truth logic for what features of the cell image the prediction model should rely on, we introduce an artificial feature to the dataset using the similar procedure to that of Kim et al. (2022). Here, we select a circular feature to demonstrate this verification process, as visualized in in the top panel of **Fig**. 3 (B), but the artificial feature can be customized to more specific applications. We then compute the average intersection-over-union score, which is typically used as a performance metric for object segmentation tasks, between the GradCAM saliency map and the ground truth averaged over the test set. We found that this metric is low (0.192) because GradCAM tended to highlight corners of the artificial feature and the highlights itself are often distorted compared to the ground truth, as shown in the middle panel of **Fig**. 3 (B). Thus, in the context of Assumption 2, we observed that GradCAM generally identifies the right hypotheses by highlighting the correct region but one should also be cautious because the highlights are not always precise. In **Supplementary Note** C, we discuss this evaluation in more detail.

### Gene Expression Data Analysis

Deep learning methods have been applied on a variety of gene expression analysis tasks, including cell type annotation, phenotype prediction and disease diagnosis. Taking advantage of well-established knowledge about gene regulatory networks, previous methods have constructed by-design models by incorporating such biological knowledge into the architecture of the neural networks. We demonstrate how the verification step can be adapted to by-design IML methods using the example of Knowledge-Primed Neural Network (KPNN) (Fortelny and Bock, 2020), which predicts the phenotypic cell state from single-cell RNA-seq (scRNA-seq) data by incorporating experimentally determined signaling pathways. The authors produce explanations of the model by inspecting the learned weights corresponding to each portion of the pathway. We constructed synthetic datasets with self-designed pathways and check whether hypotheses derived from the explanation matches the designed rules. As shown in the top panel of **Fig**. 3 (C), we designed a pathway (highlighted in red color) where two genes (0) and (1) interact with each other to control a downstream regulatory molecule (A1), which leads to the change of cell state (O). Here, we expect KPNN to assign this pathway high weights. We apply KPNN on the synthetic data and observed that all viable regulatory molecules from the related genes are assigned some importance in addition to the true pathway. We also observed that the signs of edge weights and node weights have no correlation with the actual active or repressive regulatory relationship. Specifically, only 18% edge weights in the trained model have the same signs as the designed pathway as shown in the middle panel of **Fig**. 3 (C). This analysis suggests that proposing new hypotheses by explaining a biologically informed neural network via edge weights is not guaranteed to capture the true biological mechanisms, leading to violations of Assumption 2. We also check that this finding is consistent by varying the random seeds in model training, since a different learned model will induce a different explanation for by-design models. We found that the most important edges are consistent across varied hyperparameters and random seeds as shown in **Fig**. S4. We provide remaining experimental details in **Supplementary Note** D.

## Discussion

The general workflow for knowledge discovery that we outlined above serves as an important, first start towards establishing more rigorous guidelines for IML usage in computational biology. However, the workflow can be extended for the diverse set of applications in computational biology, suggesting natural opportunities for collaboration between the IML/ML and computational biology communities.

### Additional verification steps

In this work, we focused on fundamental verification steps for IML methods to address two important assumptions made when performing knowledge discovery. It would be important for ML and computational biology researchers to work together to identify additional important steps. For example, many computational biology applications involve training a multi-modal prediction models (e.g., Srivastava et al. (2021) incorporates DNA sequence with epigenetic signals to predict TF binding and Chen et al. (2020) leverages both cancer histology images and genomic features for survival prediction). An open research question is to define a verification step to check whether explanations properly attributes importance scores to each modality. More generally, additional verification steps will help characterize potential limitations and interpretations of the IML method that is being applied.

### Improving explanation presentation

The visualizations of the explanation should be tailored to computational biology research and specialized to data types common in computational biology. One work in this direction is TF-modisco (Shrikumar et al., 2018), which synthesizes feature importance on the base-pair level into importance scores on motif level, making it easier to understand which motifs are important to the model prediction.

### Guidance on selecting IML methods

The IML community has spent a lot of effort taxonomizing explanations, but not enough work demonstrating their usefulness of downstream applications (Chen et al., 2022b). While in this work we do not explicitly provide guidance on the method selection process, except that the verification steps help to discard ineffective explanation candidates, we believe that the computation biology community would greatly benefit from a taxonomy that demonstrates what kinds of methods are useful for what kinds of applications, which requires both knowledge about IML as well as the diverse biological applications.

## Conclusion

As IML methods are increasingly used in computational biology applications for knowledge discovery, there is a need for a standardized workflow on best practices. In this work, we introduced a workflow applicable to the broad range of IML methods to address assumptions that are oft-made when using IML methods for knowledge discovery. This workflow consists of verification steps, some of which are already found in the computational biology literature (**Model Verification** and **Knowledge Verification** and a new one adapted from the IML literature (**Explanation Verification**), to enhance the reliability and generalizability of proposed hypotheses from applying IML methods to complex prediction models. We demonstrated on three different applications how **Explanation Verification** can be utilized to easily identify potential limitations of explanations.

However, we believe that this workflow marks only the beginning of a set of contributions to the foundations of IML use in computational biology. There is still a great need for joint work in the IML and computational biology communities to increasingly improve the ways in which the verification steps can be specialized for the diverse biological applications of interest. Through these collaborations, we hope that new IML problems are formulated in a way that is expected to significantly facilitate hypothesis generation and new discoveries for a wide variety of biological and biomedical contexts.

## Acknowledgements

This work was supported in part by the National Institutes of Health Common Fund 4D Nucleome Program grant UM1HG011593 (J.M.), National Institutes of Health Common Fund Cellular Senescence Network Program grant UG3CA268202 (J.M.), National Institutes of Health grants R01HG007352 (J.M.) and R01HG012303 (J.M.), and National Science Foundation grants IIS1705121 (A.T.), IIS1838017 (A.T.), IIS2046613 (A.T.), and IIS2112471 (A.T.). J.M. was additionally supported by a Guggenheim Fellowship from the John Simon Guggenheim Memorial Foundation. A.T. was additionally supported by funding from Meta, Morgan Stanley and Amazon. Any opinions, findings and conclusions or recommendations expressed in this material are those of the author(s) and do not necessarily reflect the views of any of these funding agencies.

## Author Contributions

Conceptualization, V.C., M.Y., A.T., and J.M.; Software, V.C., M.Y., W.C., and J.S.K.; Investigation, V.C., M.Y., W.C., J.S.K., A.T., and J.M.; Writing, V.C., M.Y., A.T., and J.M.; Funding Acquisition, A.T. and J.M.

## Footnotes

* Even in settings where one could perform **Knowledge Verification**, this step can be performed first to gain a better understanding of potential limitations of the IML method.

