## Supplemental Information for "Best Practices for Interpretable Machine Learning in Computational Biology"

### A Additional information on existing verification steps

For each application discussed in the **Applying Explanation Verification** section of the main text, we will also describe the ways in which the original publications (Avsec et al., 2021; Nagao et al., 2020; Fortelny and Bock, 2020) and other related work have performed **Model Verification** and **Knowledge Verification**.

#### *A.1 TF Binding*

**Model Verification:** Avsec et al. (2021) conducted model verification on BPNet by utilizing a held-out validation set. The authors partitioned the chromosomes into training, tuning, and testing sets, where chromosomes 2, 3 and 4 are used for tuning, chromosomes 1, 8 and 9 are held out for testing, and the rest are for model training. After the hyperparameters are optimized on the tuning set, the model is applied on the testing set to evaluate the ability to generalize to unseen data. Another widely adopted approach for model verification of DNA sequence-based data is *in silico mutagenesis* as demonstrated by Zhou and Troyanskaya (2015) for the DeepSEA model. DeepSEA is a CNN-based deep learning model that predicts TF binding and other chromatin features from DNA sequence and prioritizes the functional noncoding variants. The model achieves high prediction accuracy on the holdout genomic sequences (median AUC = 0.958). In addition to performance evaluations on the test sets, the authors performed model verification on DeepSEA using *in silico mutagenesis*. Experimentally validated single-nucleotide mutations (SNPs) were introduced into the input sequences and the predictions of these input sequences were compared with experimentally measured effects. For example, the authors observed that by inserting the breast cancer risk locus SNP rs4784227, the predicted binding affinity of FOXA1, increases the well-known breast cancer regulator, which matches the experimental results in Zhang et al. (2012). This verification step provides substantial evidence that the model captures the underlying rules of the prediction and generalizes well to unseen data.

**Knowledge Verification:** To generate hypotheses for knowledge verification, Avsec et al. (2021) trained BPNet to predict base-resolution binding for four transcription factors (Oct4, Sox2, Nanog and Klf4) and applied DeepLIFT to derive the base-level resolution importance scores. TF-modisco (Shrikumar et al., 2018) was then used to systematically summarize the sequence patterns and discover the important motifs. As a result, 11 representative motifs robustly discovered by running TF-modisco on different BPNet models were selected, which matched the well-characterized Oct4-Sox2, Sox2 and Klf4 motifs.

#### A.2 Imaging Data Classification

**Model Verification:** Nagao et al. (2020) performed model verification using held-out test sets, where the authors split the preprocessed cell images into training and testing sets with the ratio 8:2. Besides such partitioning of one dataset, one can also use external data to assess the generalizability of image models. One example use of external data was conducted by Coudray et al. (2018) on the DeepPATH model, a CNN trained on the well-established cancer whole-slide image dataset from The Cancer Genome Atlas (TCGA) to predict lung cancer based on histopathology images. The authors collected an independent dataset of lung cancer whole-slide images: since datasets were generated by different labs, they may contain different experimental artifacts. On these independent datasets, DeepPATH reached satisfactory classification accuracy, verifying that the model captures the important histopathological features of lung cancer.

**Knowledge Verification:** To pose hypotheses for knowledge verification, Nagao et al. (2020) used GradCAM (Selvaraju et al., 2017) to highlight important cell features. The authors found that GradCAM highlighted the area and intensity of staining for that was important for distinguishing between G1/S phase with G2 phase in cell cycle. This finding matched prior knowledge that DNA content doubles during S phase and the Golgi apparatus spreads around during the G2 phase. The results verified that the model is potentially useful for uncovering the cellular phenotypes during cell cycle. Interestingly, the authors identified patterns of the microtubule that have not been reported previously, indicating that the model could be used for new biological knowledge discovery.

#### A.3 Gene Expression

**Model Verification:** To perform model verification, KPNN (Fortelny and Bock, 2020) was evaluated on five different scRNA-seq datasets, where each dataset was modeled using a distinct neural network architecture. Similar to the previous two examples, each dataset was divided training and testing sets to avoid overfitting and evaluate each KPNN's performance.

**Knowledge Verification:** KPNN (Fortelny and Bock, 2020) is applied on the scRNA-seq data measuring cellular response to T cell receptor (TCR) and is used to propose the key regulators in the signaling pathway for knowledge verification. The network is constructed by connecting the gene expression (input nodes) with the TCR simulation (output node) via the hidden nodes representing TFs and signaling proteins, where the edges represent protein signaling and gene-regulatory interactions. The model achieves high performance on predicting the TCR based on gene expression (median AUC 0.982). To identify the key regulators in the network, the authors calculated the node weights based on the edge weights and connectivity of the nodes. They successfully identified key regulators of the signaling pathway such as NF-$\kappa$B1:RELA transcription factor complex and p38$\alpha$ MAP kinase MAPK12, which are validated by previous experimental findings.

### B TF Binding Experiment Details

This section describes the experiment details for how to perform the **Explanation Verification** step for the TF binding site prediction task. The task is to predict the TF binding sites from DNA sequences and identify the corresponding binding motifs. We adapt the workflow of the genomic experiment in Shrikumar et al. (2017): specifically, we construct a 1D CNN model to predict TF binding and leverage DeepLIFT to extract the important sequence patterns.

#### B.1 Data

To construct the dataset, we generate input DNA sequences of length 200, which consists of background noise and real motifs. The background is simulated by generating by random sampling base pairs A/T/C/G with probability 0.2/0.2/0.3/0.3. This process mimics the natural occurrences of the base pairs. The motifs are sampled from the same database as in Kheradpour and Kellis (2014), and added these motifs on non-overlapping position. The number of each kind of motif to be inserted into a sequence was determined by sampling from a Poisson distribution that was truncated to have a minimum of 1 and a maximum of 3 (when a number was sampled outside the range, it was resampled until a number within the range was achieved). For sequences containing either only one motif, the mean of the Poisson distribution was 3, and for sequences containing two motifs, the mean of the Poisson for each motif was 1.5. Each input sequence has a corresponding ground truth output that is a binary variable $\{1, 0\}$, where 1 denotes the existence of target motif(s) and 0 denotes the absence of target motif(s). Half of the dataset will have label 1 and the other half label 0. We studied three different settings: (1) if a single motif is present then label is 1, (2) if two motifs are present then the label is 1 (the cooperative condition), and (3) if one of two motifs are present then the label is 1 (the competitive condition). We generate a training dataset of size 40,000 and a test dataset size of 800.

#### B.2 Model

The model architecture is the same CNN model from Shrikumar et al. (2017)'s experiments:

1. Convolution with 50 filters of width 11
2. ReLU
3. Convolution with 50 filters of width 11
4. ReLU
5. Global Average Pooling
6. Fully connected layer with 50 neurons
7. ReLU
8. Dropout layer with probability 0.5 of leaving out units
9. Fully connected layer with 3 neurons
10. Sigmoid output

To train the model, we use the same hyperparameters as the original paper: an Adam optimizer with learning rate 0.0001, batch size 32, for 100 epochs, trained with binary cross entropy loss. We modify the output layer of the original model from 3 neurons (which predicted whether GATA, TAL1, or both motifs were present in a given sequence) to 1 neuron because we are predicting a binary output.

#### B.3 Explanation

We use DeepLIFT (Shrikumar et al., 2017), one of the most popular explanation tools, as the candidate of verification. All DeepLIFT scores are collected with "rescale on conv layers, reveal cancel on fully-connected" setting. This was found in the Deeplift paper to have the best performance. DeepLIFT provides feature importance scores for each *base pair*. To calculate the feature importance for a *motif*, we average the computed DeepLIFT score for all of the base pairs in the motif.

#### *B.4 Explanation verification*

First, we evaluate whether DeepLIFT feature importance scores suggest correct hypotheses for sequences containing only single motifs. The set of motifs we considered included SRF_disc1, AP1_disc1, GATA_disc1, TAL1_known1, IRF_known1. The logic that we want the model to learn here is simple: if the target motif is present, then predict label 1. **Fig.** S1 (Top) shows an example where all occurrences of the target motif has significant feature importance. However, **Fig.** S1 (Middle) and (Bottom) illustrate failure modes of DeepLIFT. Averaged individually across randomly generated sequences, the percentage of the following motifs falling within one standard deviation of the background noise's importance (which is centered around 0 and small in value) was SRF_disc1: 0.826, AP_disc1: 0.554, GATA_disc1: 0.434, TAL1_known1: 0.355, IRF_known1: 0.513.

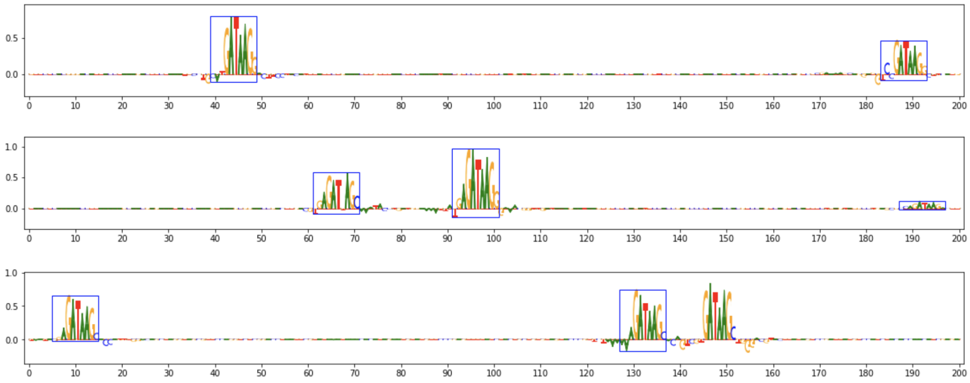


**Figure S1:** Example feature importance generated for a single motif from our synthetic dataset, where the y-axis is the feature importance score. (Top) Both occurrences of the target motif have significant feature importance. (Middle) One occurrence of the target motif has very small feature importance, which is undesirable. (Bottom) A set of base pairs not belonging to a known motif has high feature importance due to its similarities to the target motifs.

Next, we consider the hypotheses suggested by DeepLIFT in the context of more complex settings of cooperative and competitive binding (Ibarra et al., 2020). We simulate cooperative binding activities by an `AND` operator and competitive binding activities by an `OR` operator between two given motifs. The two target motifs were selected from the same list as before. We again averaged the scores across 200 randomly generated sequences for each binding setting. To measure whether how consistently DeepLIFT would identify the correct set of motifs, we measured the average scores of the base pairs that were part of the target motif against the average scores attributed to background noise and averaged this value across the entire test set. In **Table** S1 and **Table** S2, we show the percentage of average motif scores that fall within 1 standard deviation of the background score for both cooperative and competitive binding settings respectively. We find that in the cooperative binding setting, it is much more likely that the motif scores will fall in 1 standard deviation of the background score and that this occurs for multiple motif-pairs. For the competitive binding setting, however, we find that the computed DeepLIFT feature importance scores do not always reflect the expected feature importance scores. This means hypotheses generated by DeepLIFT would be inconsistent if using DeepLIFT for performing knowledge discovery.

### C Imaging Data Experiment Details

This section describes the experiment details for how to perform the **Explanation Verification** step for the imaging data classification task. The task is to predict the cell cycle phase using fluorescence microscope

**Table S1:** Percentage of motif score within 1-std of background score in the cooperative binding setting. Rows are target motif, columns are cooperative motif.

| | SRF_disc1 | AP1_disc1 | GATA_disc1 | TAL1_known1 | IRF_known1 |
|---|---|---|---|---|---|
| SRF_disc1 | | 0.51% | 0.52% | 1.14% | 0.66% |
| AP1_disc1 | 1.31% | | 24.94% | 2.62% | 6.02% |
| GATA_disc1 | 2.12% | 30.29% | | 7.64% | 45.15% |
| TAL1_known1 | 1.16% | 1.07% | 20.89% | | 2.53% |
| IRF_known1 | 0.78% | 0.27% | 99.07% | 2.27% | |

**Table S2:** Percentage of motif score within 1-std of background score in the competitive binding setting. Rows are target motif, columns are cooperative motif.

| | SRF_disc1 | AP1_disc1 | GATA_disc1 | TAL1_known1 | IRF_known1 |
|---|---|---|---|---|---|
| SRF_disc1 | | 0.13% | 0.26% | 0.64% | 0.27% |
| AP1_disc1 | 1.83% | | 0.76% | 3.67% | 2.28% |
| GATA_disc1 | 2.79% | 2.22% | | 3.29% | 3.98% |
| TAL1_known1 | 1.03% | 0.67% | 1.21% | | 1.73% |
| IRF_known1 | 1.04% | 1.08% | 0.40% | 1.01% | |

images. We train a 2D CNN model as the classifier and apply GradCAM (Selvaraju et al., 2017) on an augmented version of the dataset.

#### C.1 Data

We start with the same dataset as Nagao et al. (2020) and apply the same processing steps to the images. The images are size 128 by 128 pixels and originally have 3 channels but the authors shows that the first channel, which is Hoechst staining for DNA, is sufficient to achieve high prediction performance (90% accuracy). We then use SMERF (Kim et al., 2022) to generate the full dataset for the verification step. SMERF systematically adds artificial features to images to control the model reasoning. We chose a circular feature to demonstrate this verification step. The feature has a radius of 13 and it is put onto a random location with probability 0.5.

#### C.2 Model

We use the same CNN model architecture as in Nagao et al. (2020):

- Conv 3x3
- ReLU
- MaxPooling 2x2
- Repeat for hidden layers:
  - Conv 3x3
  - ReLU
  - MaxPooling 2x2
- Fully connected layer
- Dropout
- Fully connected layer with 2 neurons

The model is trained using the RMSprop optimizer with binary cross entropy loss. Other parameters (including the dropout rate, the number of neurons, batch size, and number of hidden layers) were selected using Bayesian optimization as in (Nagao et al., 2020).

#### C.3 Explanation

We used the GradCAM implementation from SMERF (Kim et al., 2022). GradCAM gives a feature importance for each pixel.

#### C.4 Explanation verification

We generate hypotheses using the most salient features of the GradCAM saliency map and evaluate its correctness by comparing each hypothesis to the ground truth location for where the artificial feature was added. To quantitatively measure the overlap between the ground truth location of the artificial feature and the feature identified by the saliency map, we use the intersection over union (IOU) metric. IOU calculated: first blur saliency map, take top $k$ pixels from saliency map ($k$ = number of pixels from the artificial feature, which varies by the degree to which the artificial feature is present in the image), then compute IOU. In **Fig.** S2, we show an example low versus high IOU. We find that the average IOU across the test set is 0.192 and the standard deviation is 0.166. The high standard deviation value for this simple setting shows that GradCAM does not consistently generate correct hypotheses, suggesting care when interpreting saliency maps for knowledge discovery. We refer the reader to SMERF (Kim et al., 2022) for examples of more complex settings.

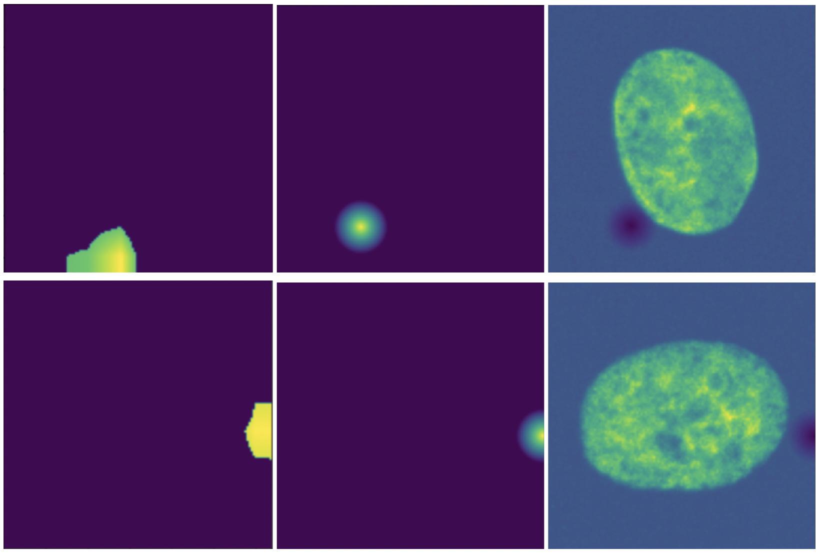

**Figure S2:** Examples of GradCAM saliency map (Left), ground truth (middle) and original image (right). The (top) row is an example with low IOU and the (bottom) row is an example with high IOU.

### D Gene Expression Experiment Details

This section describes the experiment details for how to perform the **Explanation Verification** step for the gene expression analysis task. The task is to predict the phenotypic cell state based on gene expression profile measured by single-cell RNA-seq. To demonstrate the importance of explanation verification for by-design models, we apply Knowledge-Primed Neural Network (KPNN) (Fortelny and Bock, 2020) on simulated data with simple designed pathways. We utilize the edge weights to hypothesize important pathways used by the model to make predictions.

#### D.1 Data

We construct a dataset of simulated gene expression level for $m = 100$ genes and $n = 10000$ samples, and we design the pathway in three cases with different assumptions. The $m$ nodes are divided into two groups, related genes and unrelated genes. For unrelated genes, we simulate the gene expression level by Poisson distribution with parameter $\lambda = 2$. For related genes, we assume that the genes have a positive effect on the outcome, and we use Poisson distribution with $\lambda = 5$ for positive outcomes and $\lambda = 1$ for negative outcomes.

For signaling pathway design, we assume that we know the ground truth, i.e., by which pathway the genes are affecting the phenotype. We consider the following three cases of pathway design: (1) One gene affects the outcome through a single pathway. (2) One gene affects the outcome through a single pathway, but there exist other pathways connecting the gene to the outcome, which corresponds to the redundancy discussed in the paper. (3) Two genes that are co-regulated by a hidden node in the first hidden layer jointly affect the outcome, but there exist other pathways connecting each node to the outcome.

#### D.2 Model

We use the KPNN model from Fortelny and Bock (2020). KPNN is a partially connected feed-forward neural network with biological knowledge, where the network architecture reflects the regulatory networks. Each hidden node corresponds to some proteins or genes and each edge represents some regulatory relationship along a pathway. Thus, the model architecture varies across different datasets. For the three simulation datasets, we build the KPNN based on the pathways and visualize the neural architecture on the left panel of **Fig.** S3. Specifically, the model architecture is as follows:

- Input layer with 100 gene nodes
- Partially connected layer with 4 neurons reflecting the first level of regulation
- Sigmoid
- Partially connected layer with 2 neurons reflecting the second level of regulation (only in Case 3)
- Sigmoid (Only in Case 3)
- Output layer

#### D.3 Explanation

To derive explanations from the KPNN, we interpret the edge weights as a measure of the importance of a pathway. We calculate the background edge weights using the average of all edge weights on the same level of hidden layer. We select important edges as ones with edge weights at least three times higher than the background. Note that the original paper additionally defined node weights based on the edge weights to capture the global importance of the nodes. Since we only work with one or two predictive genes and up to six regulatory elements in the simulation data, we believe that edge weights alone are sufficient to highlight the important pathways.

#### D.4 Explanation verification

To perform explanation verification, we train the designed KPNN with simulated gene expression profile and compare the hypothesized pathway based on the important edges with the ground truth pathway. The results are shown on the right panel of **Fig.** S3. The identified important edges are colored based on the

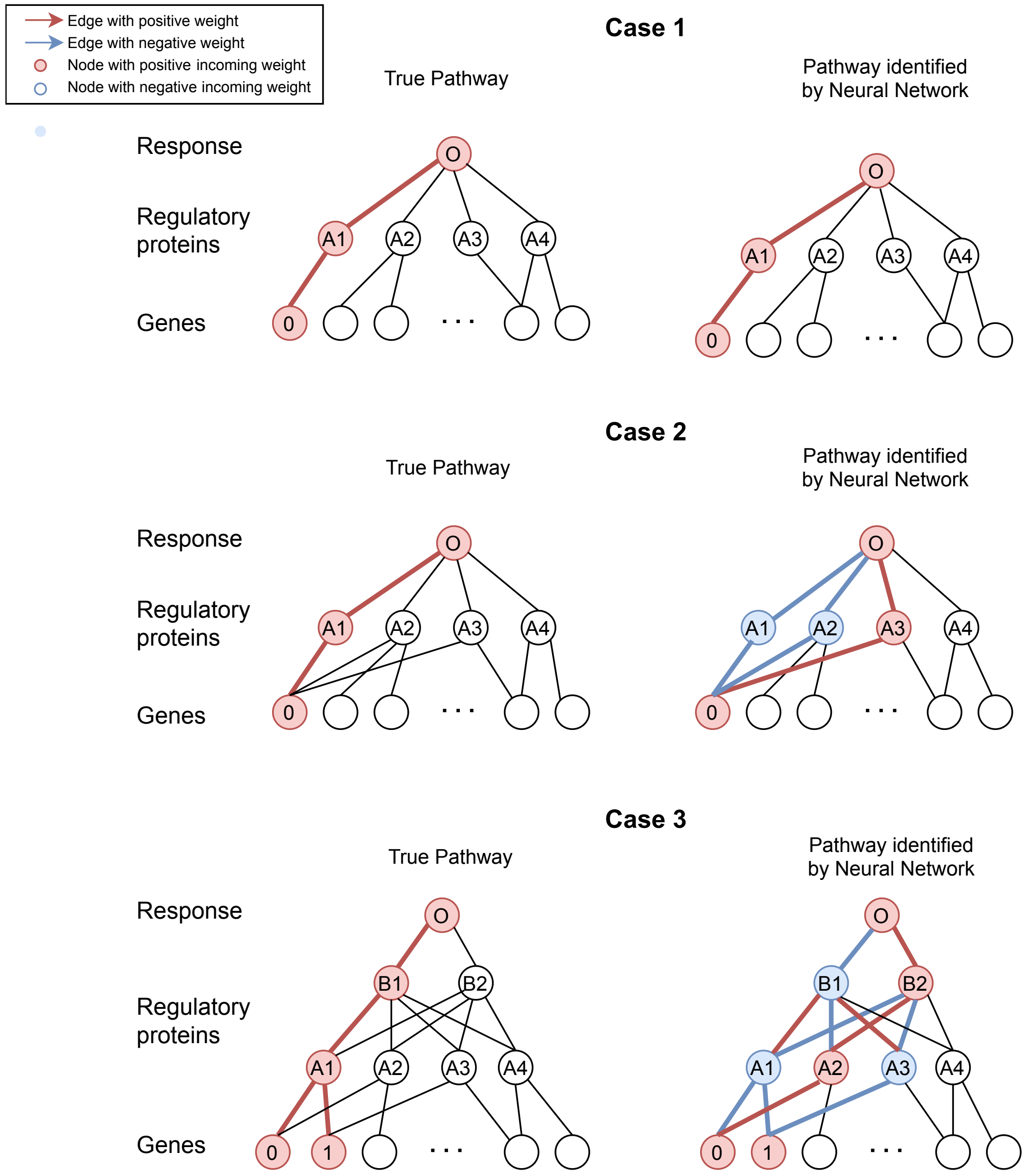


**Figure S3:** Comparison of the designed pathways and the pathways identified by the neural network. The red colors indicate strong positive edge/node weights, the blue color indicate strong negative edge/node weights, and the black colors indicate weak edge/node weights. The three cases have different pathway design. (Top) One gene affects the outcome through a single pathway. (Middle) One gene affects the outcome through a single pathway, but there exist other pathways connecting the gene to the outcome. (Bottom) Two genes that are co-regulated by a hidden node in the first hidden layer jointly affect the outcome, but there exist other pathways connecting each node to the outcome.

sign of the edge weights, where positive edges are colored in red and negative edges are colored in blue. The hidden nodes are colored based on the sign of the weights of the incoming edges.

In Case (1), the model perfectly recovers the designed true pathway. In Case (2), although the model assigns high edge weights to the true pathway, it also highlights all the other redundant pathways connecting the predictive gene to the output node. Another implication of the result is that proposing hypotheses about pathways using the sign of the intermediate edges may not be indicative of the actual regulatory relationship. The pathway with two negative edge weights, such as $G0$ -> $A1$ -> $O$, would be equivalent to the same pathway with two positive edge weights, given that there is no other incoming nodes for the intermediate hidden nodes. In Case (3), we assume that the co-regulation of gene 0 and gene 1 is important for the prediction. However, all available paths between the two predictive nodes and the

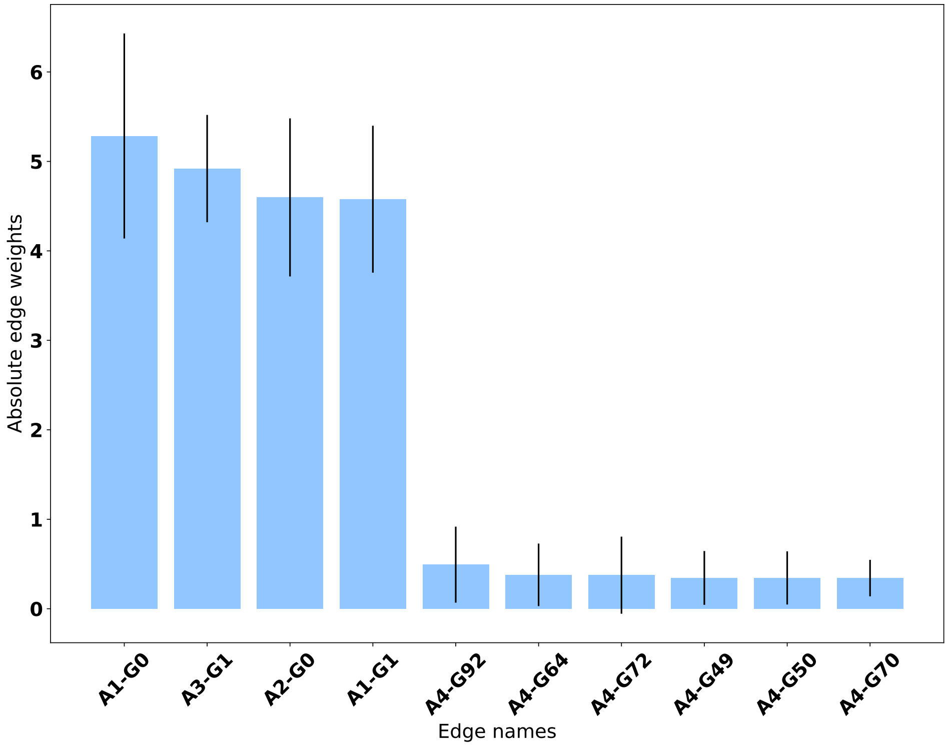


**Figure S4:** Stability check for KPNN. The bar plots display the average absolute importance scores across the different model runs.

output nodes are assigned significantly high weights. One possible explanation for the observation is that since the values of the intermediate regulatory nodes are not provided as part of the model input, it would be hard for the neural network to capture the complex regulatory relationship of a true biological system. To make sure that the explanations are consistent with the biological knowledge, additional verification steps and more advanced interpretation methods are necessary. The original paper of KPNN addressed the issues by introducing dropouts and revising the node weight metrics, which drastically improved the set of pathways identified by the model.

To further test the stability of this explanation approach, we rerun the same model on the same simulation dataset (Case 3) with different random seeds and hyperparameters such as learning rate. We visualize the edge weights ranked by the average absolute weights in **Fig.** S4. The stability test reveals that the model consistently picks the same set of important weights across the different runs, although these edges may not reflect the ground truth pathways, as discussed in the previous paragraph.
